# Stable and assayable polymicrobial human airway model reveals complex interactions between *Pseudomonas aeruginosa* and lung commensals

**DOI:** 10.1101/2025.10.30.685207

**Authors:** Sarah M. Spencer, Karla N. Valenzuela-Valderas, Brendan M. Leung

## Abstract

Polymicrobial-host crosstalk shapes airway barrier integrity and inflammation, yet standard *in vitro* systems rarely sustain interaction-dependent phenotypes. We engineered an aqueous two-phase system (ATPS) confined bronchial co-culture that stabilized day-scale assays while maintaining epithelial function. A 16HBE14o-/HUVEC cell insert model was challenged with *Pseudomonas aeruginosa* PA01, *Streptococcus pneumoniae* D39, and the commensals *Rothia mucilaginosa* and *Lactobacillus casei* in mono- and polymicrobial combinations. We evaluated epithelium permeability to FITC-dextran, cell junction integrity, bacterial-induced cytotoxicity, IL-6 and IL-8 release and bacterial viability. ATPS preserved a workable assay window and bacterial confinement over 24h. PA01 disrupted barrier integrity, and commensals mitigated this pathogenic effect, whereas PA01 co-cultured with *S. pneumoniae* exhibited synergistic damage effects on the lung epithelium. Junctional imaging corroborated functional readouts, and cytotoxicity remained low across conditions. Cytokine shifts were condition-specific but modest, demonstrating compatibility for soluble mediator profiling. We generated a human airway-microbiome model in which ATPS confinement enabled modelling of dynamic lung-microbiome interactions, reproducing known *P. aeruginosa* pathogenic effects and commensal protection of the epithelium.

## 1. Introduction

The human airway microbiome dynamically shapes respiratory health [1]. Airway pathophysiology arises from complex interaction-dependent dynamics between multiple species and the host [2]. Polymicrobial communities shape barrier integrity, antimicrobial defence, and the epithelial inflammatory landscape [3], [4]. Failing to consider polymicrobial interactions in the respiratory system obscures the clinical relevance of current host-microbe studies in healthy and diseased models.

In cystic fibrosis, impaired mucociliary clearance and thickened mucus create niches that foster polymicrobial communities including *Pseudomonas aeruginosa*, *Staphylococcus aureus*, and *Burkholderia* [5]. Polymicrobial interactions between these taxa markedly alter microbial behaviour. For example, *Staphylococcus* can enhance *P. aeruginosa* persistence and antibiotic tolerance, mediating shifts in microbiome composition [6]. Similar polymicrobial dynamics are also observed in chronic obstructive pulmonary disease (COPD) and bronchiectasis, where exacerbation risk and lung function decline correlate with changes in microbiome composition [7]. Even in lung cancer, the airway microbiome has been linked to tumor progression and therapeutic response [7]. Across these conditions, understanding polymicrobial interactions is essential for explaining pathophysiology and guiding effective interventions. Despite this, most *in vitro* models remain single-species due to challenges in co-culturing host and microbial communities.

Microbe-mammalian co-cultures often collapse due to bacterial overgrowth, nutrient depletion, or epithelial cytotoxicity [8]. These models typically yield only endpoint measurements over short assay windows, missing dynamic emergent interactions. Animal models capture polymicrobial complexity but differ from human physiology and have limited mechanistic and assay resolution. We therefore aimed to generate a model geometry that bridges the gap between single-species studies and animal models, preserving epithelial viability while revealing polymicrobial influences over day-scale readouts.

Spatially confining bacteria can be achieved using the PEG-DEX <u>a</u>queous two-<u>p</u>hase <u>s</u>ystem (ATPS), which forms an immiscible microdroplet that confines bacteria, limiting bulk mixing and thus bacterial overgrowth, while permitting the diffusible exchange of microbial and host-derived factors [9]. We previously showed that bacterial confinement creates a localized, highly effective area of infection at the droplet-epithelium interface, thereby accelerating the detection of interaction-dependent effects [10]. ATPS confinement prevents collapse of the co-culture, extending the assay window to the day scale and enabling resolution of interaction-dependent effects that are typically obscured in conventional liquid co-culture.

Here, we developed an ATPS-confined microbial-bronchial epithelial airway model to recapitulate interaction-dependent airway phenotypes, including *P. aeruginosa*-driven barrier disruption and inflammatory signalling, while also revealing attenuation by commensal taxa. Our model preserves epithelial function over day-scale exposure to bacteria while reproducing known pathogenic and commensal effects on epithelium integrity and secreted factors.

## 2. Methods

### 2.1 Mammalian cell culture

Human bronchial epithelial cells (16HBE14o-) were maintained in a 1:1 DMEM and Ham’s F-12 (DMEM-F12, Gibco) mixture supplemented with 10% fetal bovine serum and 1% antibiotic-antimycotic. Human umbilical vein endothelial cells (HUVEC) were cultured in Endothelial Cell Growth Medium 2 (EGM-2, Promocell) supplemented with the accompanying supplement mix. All cell lines were cultured at 37°C in 5% CO2.

### 2.2 Bacterial strains and culture conditions

*Pseudomonas ae*ruginosa PA01 (provided by Dr. Zhenyu Cheng, Dalhousie University) was cultured on LB at 37°C overnight. *Streptococcus pneumonia* D39 (provided by Dr. Jason Leblanc, Dalhousie University) was cultured on blood agar at 37°C overnight. *Lacticaseibacillus casei* 03 (ATCC 393™) was cultured on MRS agar at 37°C for 48h, and *Rothia mucilaginosa* 5762/67 (ATCC 25296™) was cultured on BHI at 37°C overnight. Prior to experiments, overnight liquid cultures were inoculated with 1-2 colonies and incubated at 37°C with shaking at 200rpm for 15-18h. For *S. pneumoniae* cultures, 5 colonies were used to inoculate 4 mL of Todd Hewitt broth (BD 249240) supplemented with 2% Difco™ yeast extract and 10% Oxyrase® (Oxyrase inc. OB-0100) and were incubated at 37°C in 5% CO2 with shaking at 135rpm for 15-18h.

Optical density was measured at 600 nm using a Varioskan plate reader (Thermo Fisher Scientific) by transferring 100 µL of culture into a 96-well plate. Growth curves were generated to determine exponential growth periods for each strain. Sub-cultures were generated by diluting the overnight culture into fresh media and incubating for 3-4h prior to experimentation to ensure bacteria were in the exponential growth phase when introduced to the model system. This incubation time was determined from strain-specific growth curves.

### 2.3 Airway epithelial-endothelial model

Milicell® hanging cell culture inserts (12-well, 8 µm pore size, PCEP12H48, MiliporeSigma) were used as the model platform. Inserts were coated by submersion in 0.05 mg/mL type I bovine collagen (PureCol® Advanced Biomatrix, 5005-100ml) diluted in molecular grade water for 2h at RT (protocol loosely adapted from [12]). Collagen solution was aspirated, and membranes were air-dried until no longer visibly wet. Inserts were inverted, and 1 x 10^5^ HUVEC cells in 200 µL of media were seeded on the basolateral surface. Inserts were incubated in this orientation for 2h at 37°C, 5% CO2. After 2h cell culture inserts were reverted to their upright orientation and 1 x 10^5^ 16HBE cells in 500µL of media were seeded on the apical surface (adapted from [13]). The model was maintained with media containing progressively lower glucose and serum concentrations. On day 1, the model was grown in a 1:1 mixture of DMEM-F12 and EGM-2. On day 3, the media was changed to a mix of 25% DMEM-F12, 25% Low-glucose DMEM-F12 with 1% FBS, and 50% of EGM-2; at this point, antimicrobials were removed. Lastly, on day 5, a 1:1 mixture of Low-glucose DMEM-F12 with 1% FBS and EGM-2. ATPS containing bacteria were applied on day 6.

### 2.4 ATPS preparation and addition to the airway model

The aqueous two-phase system (ATPS) consisted of 5% (w/v) polyethylene glycol (PEG, 35kDa; Sigma Aldrich 8.18892.1000) and 5% (w/v) dextran (DEX 500kDa, Pharmacosmos 5510 0500 9007) dissolved in a 1:1 mixture of Low-glucose DMEM-F12 with 1% FBS and EGM-2 as previously described [9][10]. Briefly, the mixture of dissolved DEX and PEG was filter-sterilized, centrifuged at 3000g for 90 min, and the top (DEX) and bottom (PEG) phases were carefully separated. Bacterial cultures in the exponential phase were quantified by OD_600_ measurement. Cultures were diluted to the desired experimental concentrations (*P. aeruginosa*, OD_600_=0.0005*; S. pneumoniae*, OD_600_=0.2; *R. mucilaginosa*, OD_600_=0.025; *L. casei*, OD_600_=0.025). Bacteria were pelleted by centrifugation at 8000g for 5 min and resuspended in the DEX-rich phase by pipette mixing and gentle vortexing.

Before adding the bacteria to the airway model, the apical media were carefully removed and replaced with 400 µL of PEG-rich phase. A 0.5µL droplet of bacteria-containing DEX was then added to establish the ATPS, as described previously [9]. Dextran droplet deposition was confirmed with phase contrast microscopy. Models were incubated at 37°C with 5% CO_2_ for 24h. Afterward, the models were imaged by phase-contrast microscopy (4x objective; EVOS™ FL Auto 2 Imaging System) to assess bacterial growth and ATPS confinement. Inserts were then transferred to new 12-well plates. Apical media was gently extracted for ELISA and CFU quantification.

### 2.5. Dextran permeability assay

Epithelial barrier permeability was measured using FITC-dextran (40kDa; Sigma Aldrich FD40S). FITC-dextran was dissolved to 1 mg/mL in 1:1 phenol red-free DMEM-F12, supplemented with 1% FBS and 1% Anti-anti, sterile filtered through a 0.22 µm syringe filter. Then, 200 µL of FITC-dextran was added to the apical compartment of the insert, and 800 µL of phenol red-free media was added to the basolateral compartment. Basolateral fluorescence was measured at 0.25h, 1h, 3h, 6h, and 24h, while cultures were maintained at 37°C in 5% CO_2_. At each time point, 30 µL of basolateral media was collected, diluted in 270µL of PBS, and read in triplicate (3 x 100 µL) in black 96-well plates using a Varioskan plate reader (excitation/emission: 490/520nm). A 100% permeability control consisted of 800 µL of media in the bottom container and 250 µL FITC-dextran solution in the upper container. This protocol was loosely adapted from Faber and McCullough [14].

### 2.6 ELISA

Apical samples were centrifuged at 1000 g for 10 min, and supernatants were stored at -80°C until analysis. Cytokine levels were quantified using Legend max ™ and ELISA max ™ Delux Kits (BioLegend) according to the manufacturer’s protocols.

### 2.7 Live-dead cytotoxicity assay

Cytotoxicity was assessed using a Live/Dead viability assay (Calcein-AM and Ethidium Homodimer-1; Thermo Fisher Scientific, L3224) with Hoechst nuclear stain (Thermo Fisher Scientific, 33342). Staining was performed according to the manufacturer’s protocol after ATPS removal at the 24h endpoint. Fluorescence and phase-contrast images were acquired using an EVOS™ FL Auto 2 Imaging System (Thermo Fisher Scientific) and processed in Fiji/ImageJ (v2.16.0).

### 2.8 Immunofluorescence

Following permeability assays, inserts were rinsed with PBS and fixed in 4% paraformaldehyde (Thermo Fisher Scientific) at 4°C for 24-48h. The fixative was then removed, and the membranes were rinsed and stored in PBS at 4°C until staining. Membranes were permeabilized in 0.2% (w/v) Triton X-100 (Sigma-Aldrich X100) for 4 min, excised from inserts with a scalpel and transferred to 24-well plates. Samples were incubated overnight at 4°C with primary antibodies against E-Cadherin (1:200; Abcam ab40772) or ZO-1 (1:200; Abcam ab221547) diluted in 1% BSA (Sigma Aldrich A7906) in PBS. After 3 PBS washes, Alexa Fluor 647-conjugated donkey anti-rabbit IgG (1:1000; Abcam ab150075) was applied for 2h, followed by DAPI nuclear stain (1:1000; 20 min). Membranes were mounted on slides, and Z-stacks were acquired by confocal microscopy (Leica TCS SP8 laser-scanning confocal). Stacks were processed in Fiji/ImageJ using AnalyzeSkeleton [15] to obtain the number of junctions and endpoints within each image. Junctional continuity for a field was defined as the junctional fraction (Formula 1) and averaged across three fields per sample (biological replicates = 3), values were then normalized to the matched experimental control.

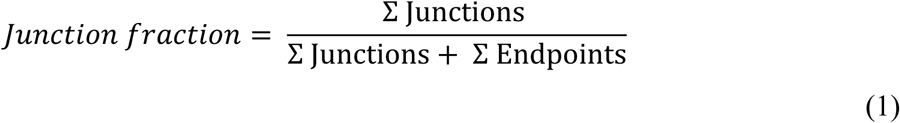

### 2.9 Colony-forming units (CFU) quantification

Apical media was collected from the model inserts and combined with one PBS wash. These samples were serially diluted (10^0^–10^8^). Dilutions (10 µL) were spot-plated on the corresponding agar. Co-cultures of PA01 and *R. mucilagenosa* were cultured on BHI agar supplemented with 10 mg/L colistin sulphate salt (Sigma Aldrich C4461) to inhibit PA01 growth. Co-cultures of PA01 and *L. casei* were plated on MRS media, which is selective for Lactobacillus. Plates were incubated for 24-48h, and colonies were enumerated to calculate CFU/mL in mono- and polymicrobial conditions.

### 2.10 Statistical Analysis

Analyses were done in R (v4.x) with {tidyverse}, {rstatix}, {lme4}/{lmerTest}, and {emmeans}. Biological replicates were independent cultures; technical replicates were averaged per sample before analysis. Normalized outcomes were tested against their benchmark (dextran permeability: PAO1 = 100%; junction continuity: control = 1.0) using two-sided one-sample *t*-tests (Wilcoxon if non-normal). Across-group comparisons at a single time point used Welch one-way ANOVA with pairwise Welch *t*-tests; multiplicity was controlled by Holm or Benjamini–Hochberg as specified in figure legends. Adjusted *p* values are reported in the results.

## 3. Results

### 3.1 Airway epithelium model maintains barrier function and junctional architecture with ATPS formulation

Our model, consisting of a bronchial epithelial-endothelial co-culture on cell culture inserts, mimics the distal airway barrier and allows us to perturb and measure barrier integrity, a clinically relevant readout (Fig.1A). Before introducing bacteria, we verified that the culture protocol provided a high-integrity baseline suitable for detecting perturbations and evaluated PEG influences on epithelium permeability. FITC-Dextran permeability was measured over 24h in untreated controls and in groups pre-exposed to the PEG-rich phase (no bacteria) for 24 h and 48h (Figure 1B). Permeability values were normalized to a no-cell insert (100% permeability). The 24h PEG group remained indistinguishable from controls across the time course with <5% permeability at 24h (mean ± SD; 2.5±2.5%, n=3 per timepoint). In groups exposed to PEG for 48h, permeability was increased in comparison to the 24h exposure group, but remained low (∼15% ± 7%) and still within error of the control. Complementary immunofluorescence showed continuous junctional E-Cadherin and Zonula Occludens-1 (ZO-1) labelling in all conditions, confirming preserved junctional architecture (Figure 1C). These results demonstrate that the ATPS formulation does not degrade baseline barrier function and that the model retains sufficient dynamic range to resolve barrier-disruptive stimuli without saturating the assay window.

**Figure 1.**
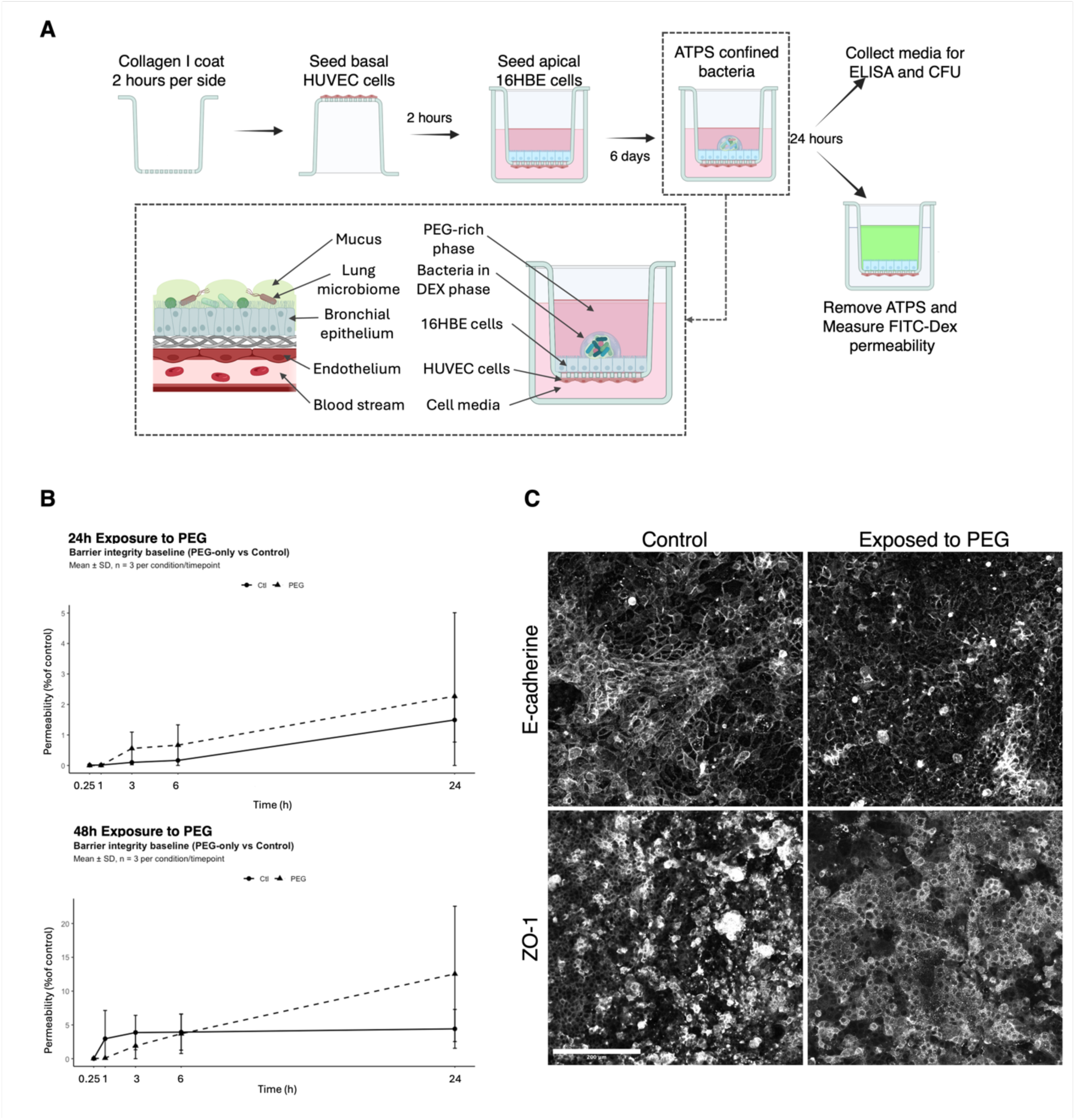
Development of an airway epithelial-endothelial model that preserves barrier integrity. **(A)** Diagram showing step-by-step preparation of the airway model, experimental readouts and similarity of its components with lung epithelium. **(B)**Baseline barrier integrity is preserved under PEG exposure. FITC–dextran permeability over 24 h (top) for airway models pre-exposed to PEG (dashed line) vs untreated controls (solid line) for 24h (top) and 48h (bottom). Points are mean ± SD (*n* = 3 per condition/timepoint). (**C**) Representative confocal images of E-cadherin and ZO-1 junction proteins showing continuous junctional localization in control and PEG-exposed models. Scale bar: 200µm. Images acquired on a Leica TCS SP8 (LIGHTNING) confocal microscope.

### 3.2 ATPS formulation sustains spatially confined mono- and polymicrobial growth for up to and over 24h

To evaluate whether the ATPS formulation used in this study would confine mono- and polymicrobial growth during the experimental duration (24h), we tracked the inoculated DEX phase of the ATPS microdroplet deposited on the PEG phase without mammalian cells for 0-45h. Across all conditions, the DEX droplet footprint and phase boundary remained intact for at least 24h, indicating sustained spatial confinement of the bacteria within the DEX phase (Figure 2). Notably, the PA01 conditions, both mono- and polymicrobial, exhibited peripheral string-like protrusions at later time points (generally showing up around 12h). As shown in the insets in Figure 2, these protrusions did not indicate a breach of the PEG/DEX interface, with a clear contrast in bacterial cell density within the droplet and no visible bacteria outside the phase boundary.

**Figure 2.**
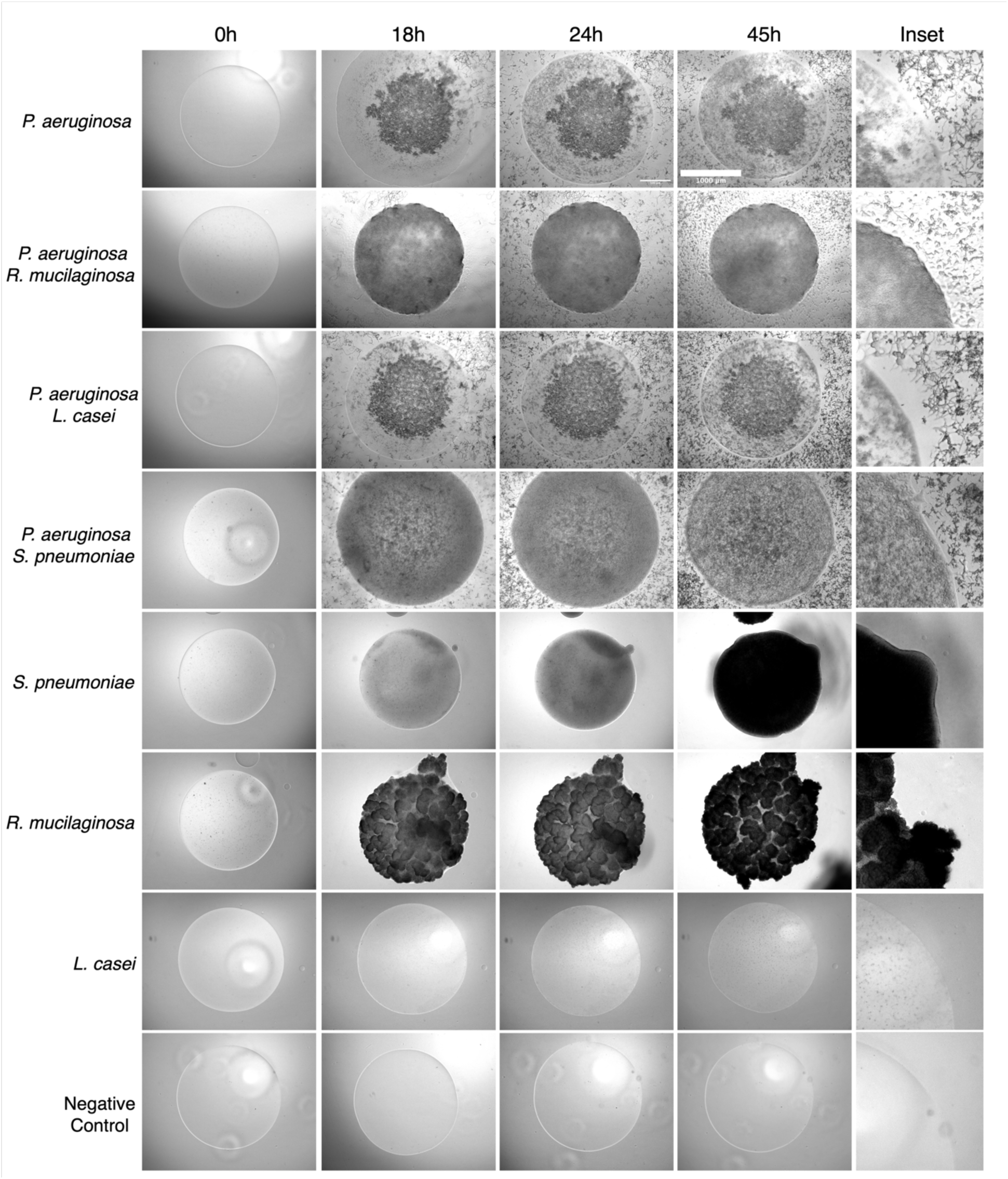
ATPS confinement enables stable, spatially confined mono- and polymicrobial growth over ≥ 24h. Phase-contrast time-lapse of Dex-phase microdroplets deposited on tissue culture plates under eight microbial conditions. Columns show 0, 18, 24, and 45 h and inset from the 45h time point. Scale bar = 1000µm. Images acquired on an EVOS™ FL Auto 2 Imaging system.

### 3.3 Without ATPS, *P. aeruginosa* rapidly collapses the epithelial barrier, preventing observation of polymicrobial interactions

To demonstrate the necessity of ATPS for observation of nuanced polymicrobial effects on epithelial health, the airway epithelial model was exposed to *P. aeruginosa* in either confined (ATPS) or unconfined (standard liquid culture) conditions. In both conditions, equivalent dosages of bacteria were used (0.5µL droplet at OD_600_=0.0005). Following inoculation, cultures were maintained for 24h under identical conditions. FITC-dextran permeability was quantified longitudinally over the following 24h. With equivalent bacterial concentration, unconfined growth of *P. aeruginosa* caused a rapid increase in airway barrier permeability that saturated the assay window within hours. In contrast, *P. aeruginosa* growth confined with ATPS remained within interpretable permeability ranges for the duration of the assay (over 24h) (Fig.3A). In the permeability assay time course measurements, unconfined *P. aeruginosa* reached ≈80% barrier loss by 6h (mean ± SEM = 81.8 ± 5.23%), while ATPS confined cultures were all <25% permeability at the same time point (12.3 ± 5.79%). By 24h, unconfined *P. aeruginosa* reached near complete membrane barrier destruction (98.3 ± 4.93%), whereas confined *P. aeruginosa* maintained low to moderate permeability (50.3 ± 8.24%). Immunofluorescence at 24h corroborated these functional results; unconfined *P. aeruginosa* caused near complete loss of the epithelial monolayer and junctional protein signal (E-cadherin, ZO-1), leaving sparse nuclear DAPI signal, whereas ATPS confined *P. aeruginosa* preserved a near continuous monolayer (Fig. 3B).

**Figure 3:**
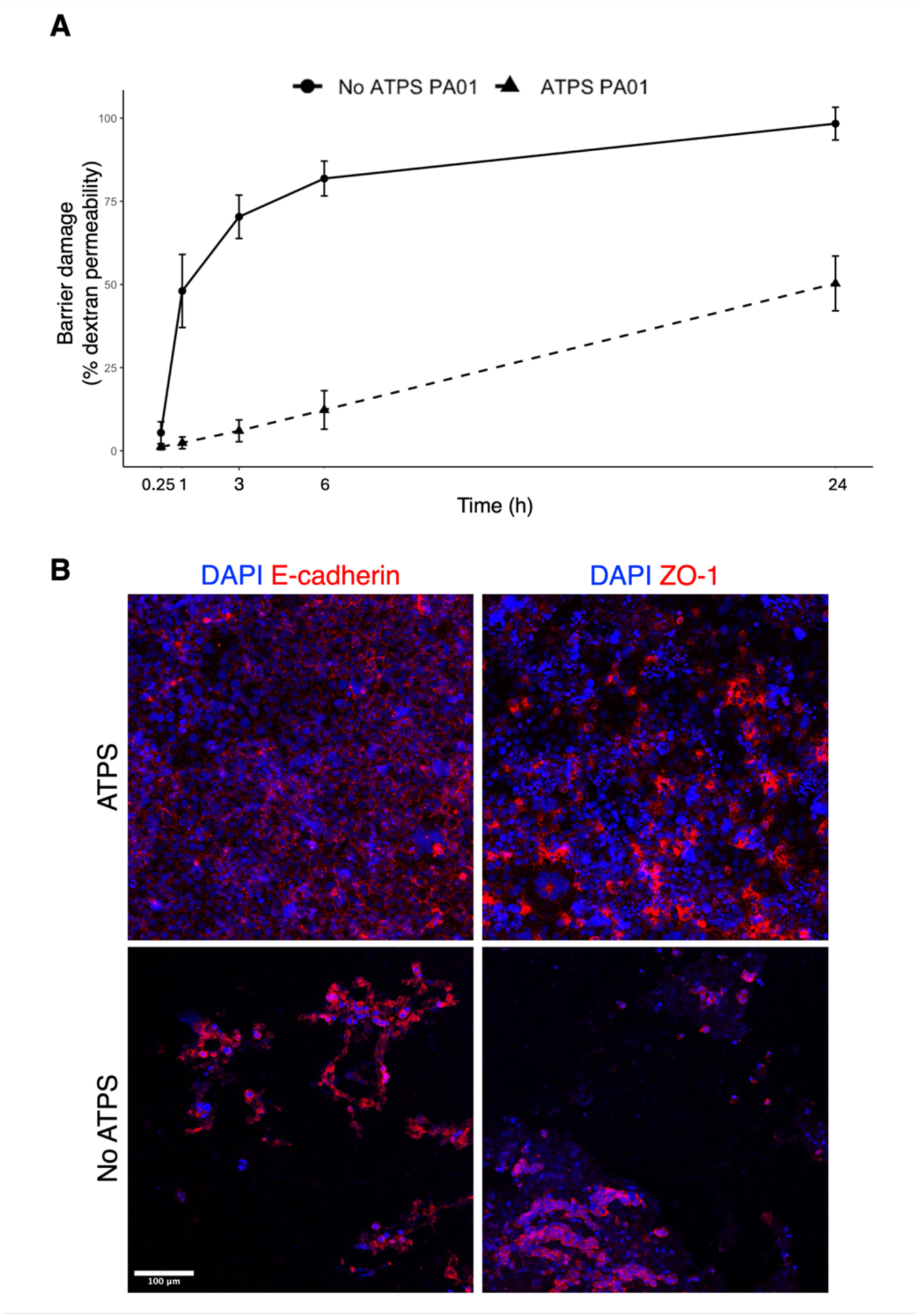
ATPS maintains tissue functionality and junctional architecture. (**A**) FITC–dextran permeability time course for the airway epithelial model exposed to *P. aeruginosa* PAO1 under unconfined (liquid or “no ATPS”) or confined (ATPS) conditions. Equivalent inocula were used (0.5 µL at OD600 = 0.0005). Cultures were maintained for 24 h prior to permeability measurements, which were collected over the following 24 h. Data points represent are mean ± SEM (n=3 biological replicates). (**B**) Representative IF images at 24 h showing nuclei (DAPI, Blue), E-cadherin and ZO-1 (Cy-5, Red). Unconfined PAO1 produced extensive epithelial denudation with loss of junctional E-cadherin/ZO-1 signal. ATPS confinement preserved a contiguous monolayer with intact E-cadherin/ZO-1 junctions. Scale bar = 100µm, images acquired on Leica TCS SP8 laser-scanning (LIGHTNING) confocal.

### 3.4 Commensals attenuate *P. aeruginosa*-mediated barrier disruption, while the addition of *S. pneumoniae* potentiates damage

Because the airway epithelium forms the first line of defence against inhaled microbes and barrier dysfunction is a hallmark of disease in asthma, CF, and COPD, we leveraged well-described lung pathogens and commensal bacteria, including the *P. aeruginosa* PA01 strain, known to cause acute lung infections [11] and the commensals *Rothia mucilaginosa* and *Lactobacillus casei*. We also used *Streptococcus pneumoniae*, commonly isolated in cystic fibrosis (CF) patients, which, in combination with *P. aeruginosa*, achieves supra-additive effects. First, to test whether commensal species mitigate pathogen-driven barrier damage on our airway model, we quantified FITC-Dextran permeability (normalized to the paired *P. aeruginosa* (PA01) = 100%, design consistent null “no change from PA01”) across the mono- and polymicrobial cultures and assessed junctional architecture using E-cadherin and ZO-1. Relative to *P. aeruginosa* alone, polymicrobial culture with either *R. mucilaginosa* or *L. casei* significantly reduced barrier damage to roughly half of the PA01 benchmark (Fig. 3A, mean barrier damage, PA01 with *R. mucilaginosa* = 50.7%, PA01 with *L. casei* = 58.3%). Commensals alone were indistinguishable from the no-bacteria control (Fig. 4A). Commensals conferred robust protection against *P. aeruginosa*-mediated functional barrier disruption in this model. Interestingly, *S. pneumoniae* alone did not increase permeability relative to control; however, *P. aeruginosa* in co-culture with *S. pneumoniae* produced super-additive injury to the model, exceeding the expected effect from the simple additive of the single pathogen conditions (Figure 3A).

**Figure 4:**
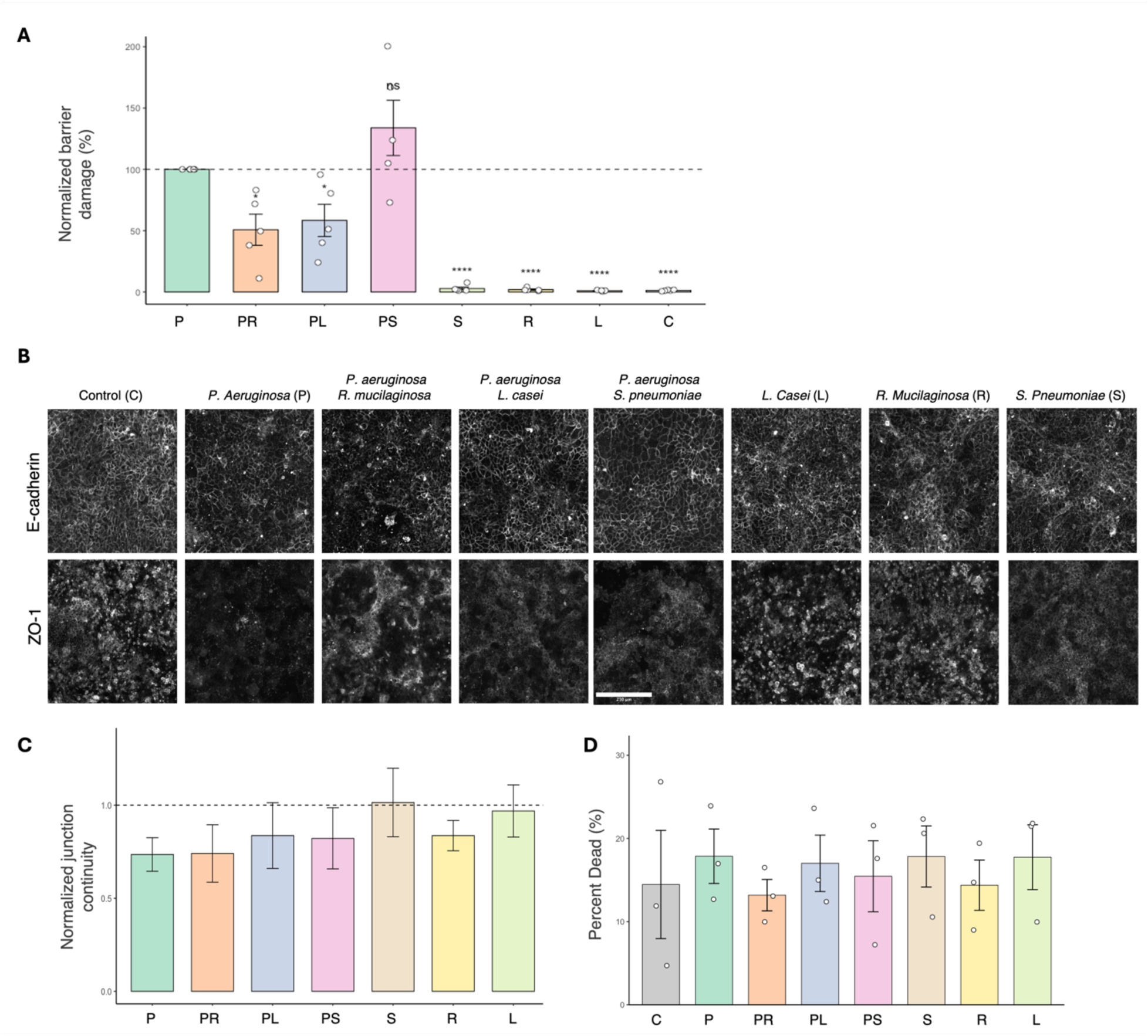
Commensals attenuate *P. aeruginosa*-mediated barrier damage while preserving junctional architecture. **(A)** Dextran-permeability readout expressed as “normalized barrier damage (%)”: for each experiment, values were scaled to the paired *P. aeruginosa* condition (*P. aeruginosa* = 100%). Bars are mean ± SEM (n=5) with overlaid points indicating biological replicates. Significance was assessed for each group against 100% using two-sided one-sample t-tests; asterisks denote adjusted *p* values after Benjamini–Hochberg FDR correction (*p* < 0.05 = *, < 0.01 = **, < 0.001 = ***, < 10⁻⁴ = ****; “ns” otherwise). *P. aeruginosa* + *R. mucilaginosa* (PR) and PAO1+*L. casei* (PL) reduces damage relative to *P. aeruginosa* (P), whereas *P. aeruginosa* +*S. pneumoniae* (PS) shows the highest mean damage. Single commensals (R, L) and S exhibit minimal damage. Control is denoted as C. **(B)** Representative confocal images of E-cadherin and ZO-1 immunofluorescence. Images illustrate junctional disruption by *P. aeruginosa* and its mitigation by commensals (PR, PL co-cultures). Scale bar, 250 μm. Images acquired with a Leica TCS SP8 confocal microscope. **(C)** Junctional continuity (E-cadherin network continuity ratio) normalized to the matched control within each experiment (C = 1; dashed line). Bars are mean ± SEM, n = 3. **(D)** Percent dead cells (live/dead staining) across groups; bars show mean ± SEM (n=3) with biological replicates overlaid. Group differences were evaluated with one-way Welch ANOVA followed by pairwise Welch *t*-tests with Holm correction; no condition exhibits significant cell death that could account for permeability changes.

Structural readouts supported the FITC-Dextran permeability results. Immunofluorescence imaging showed preservation of continuous E-cadherin and ZO-1 expression in co-cultures relative to *P. aeruginosa* alone (Fig. 4B). Quantification of E-Cadherin junction continuity (normalized to control = 1) showed similar directionality and higher continuity in commensal polymicrobial groups compared to *P. aeruginosa* alone, although this trend did not reach statistical significance under our current sampling (n=3; Fig. 4C). ZO-1 was not quantified but qualitatively mirrored the E-cadherin findings (Figure 4B).

Finally, mammalian cell viability assessment ruled out overt cytotoxicity as a confounding factor in the polymicrobial interactions. Percent dead cells were low and statistically indistinguishable across conditions (Fig. 4D), indicating that the differential barrier leakage is not attributable to condition-specific cytotoxicity or epithelial loss. The functional (FITC-Dextran permeability) and structural (junctional markers) data indicate that commensals mitigate PA01-induced barrier damage, with significance at the functional level and visual corroboration at the architectural level.

### 3.4 Differential IL-6 and IL-8 secretion

We then evaluated cytokine release to understand the airway model’s inflammatory response across mono- and polymicrobial conditions. Overall, we observed that *P. aeruginosa* exerts an immunosuppressive phenotype in the airway model that can be reversed when co-cultured with commensals (Fig. 5). PA01, grown in combination with the commensals *R. mucilaginosa* (0.908 ± 0.119 ng/mL) and *L. casei* (≈1.3x, mean ± SEM: 1.206 ± 0.174 ng/mL), increases IL-8 secretion (Fig. 5A). As expected, commensals alone show marginal IL-8 induction. Interestingly, PA01 and *S. pneumoniae* co-culture reduce IL-8 secretion levels compared with *S. pneumoniae* monoculture (≈1.5x control; mean ± SEM: 1.430 ±0.2248 ng/mL), demonstrating the strong immunosuppressive effects of *P. aeruginosa* (Fig. 5A). IL-6 secretion was also hindered by PA01 and remained lower even in the presence of commensals (P/PR/PL ≈0.5-0.9x, mean range: 5.399-8.099 ng/mL), whereas monocultures of *S. pneumoniae*, *L. casei*, and *R. mucilaginosa* remained moderately elevated (≈1.4x, 1.4x, and 1.2x, respectively). Similarly to IL-8, co-culturing *S. pneumoniae* with PA01 reduced IL-6 secretion observed in *S. pneumoniae* alone. These results demonstrate the platform’s feasibility, showing that it can recapitulate inflammatory responses to pathogens and co-culture effects.

**Figure 5:**
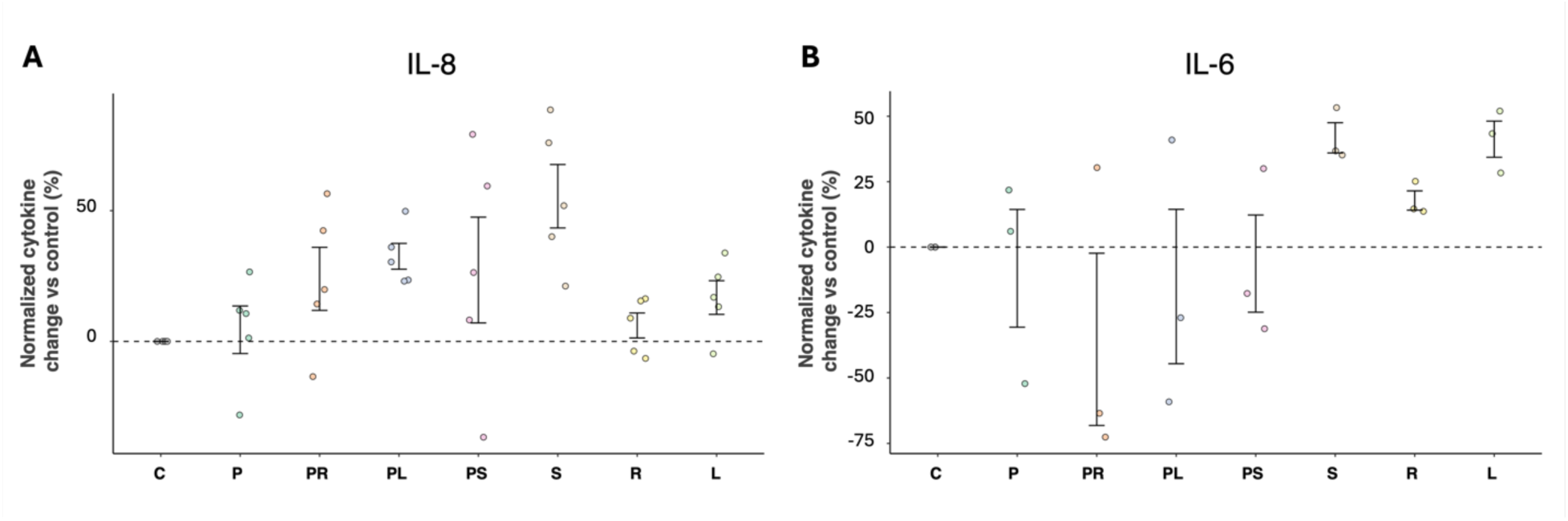
Normalized percent change in cytokine response across conditions. (**A**) IL-8 and (**B**) IL-6 measured by ELISA and summarized as percent change relative to the control condition from the same biological replicate: %Δ = 100 x (condition/control -1), control = 0% (dashed line). Each dot is a biological replicate (triplicates averaged to one point per run). Bars represent SEM. Condition abbreviations: **P** = *P. aeruginosa* PA01, **PR**/**PL**/**PS** = PA01 + **R** (*R. mucilaginosa*), **L** (*L. casei*), or **S** (*S. pneumoniae*), respectively. **R**, **L**, **S** = single species alone.

### 3.5 *P. aeruginosa* growth at 24h is not impeded by polymicrobial conditions

To rule out pathogen loss as a confounder for barrier integrity readouts, we quantified bacterial loads at the experimental endpoint. *P. aeruginosa* was inoculated into the airway model at ≈2.0 x10^6^ CFU/mL and consistently reached >1.0 x10^8^ CFU/mL across all replicates in mono- and polymicrobial conditions after 24h, indicating bacterial expansion and no evidence of commensal-mediated killing or suppression of PA01 growth. In parallel, *L. casei* CFU/mL at experimental endpoint were of similar magnitude in mono- vs polymicrobial culture (mean ± SEM, mono: 4.23 x10^5^ ± 2.00 x10^5^; poly: 3.38 x10^5^ ± 1.64 x10^5^; n=3); however, it was well below the *L. casei* inoculum (≈7.37 x 10^7^ CFU/mL). *R. mucilaginosa* (≈4.33 x 10^3^ CFU/mL starting inoculum) and *S. pneumoniae* (≈5.67 x10^7^ CFU/mL starting inoculum) were not enumerated at the experimental endpoint due to *R. mucilaginosa* generating biofilms that could not be disrupted from ATPS, and *S. pneumoniae* co-cultures were incompatible with CFU enumeration due to the lack of a selective medium.

## 4. Discussion

Polymicrobial–host interactions govern airway barrier integrity, antimicrobial defense, and inflammatory tone across diseases such as CF, COPD, bronchiectasis, and other lung conditions [7], yet most accessible in-vitro systems remain mono-microbial and collapse before emergent dynamics can be resolved. To address this limitation, we developed an airway-lung model that recapitulates microbial-host interactions. ATPS transforms a rapidly collapsing co-culture into an assayable, day-scale system suitable for longitudinal functional and structural readouts. With this approach we can tease out nuanced host-polymicrobial interactions that are otherwise obscured by the rapid system collapse caused by PA01. *R. mucilaginosa* and *L. casei* mitigate PA01 mediated damage and *S. pneumoniae* combined with PA01 trends towards synergistic epithelium damage. Viability assays reveal minimal change in viability between conditions, indicating that observed effects are not confounded by mass cytotoxicity. This study establishes an adaptable, interaction-aware airway platform that enables controlled extended duration polymicrobial co-culture with live imaging and routine endpoints to generate structural and functional readouts that expose interaction-dependent phenotypes that are generally inaccessible to standard co-cultures. Collectively, these results recapitulate interaction phenotypes and establish the platform’s predictive utility for probing less defined host-microbe and microbe-microbe interactions.

Airway barrier integrity is a clinically meaningful indicator of lung health because it represents the first line of defense against inhaled microbes and particles. Tight junctions and adherens junctions regulate paracellular permeability, and their disruption allows pathogens, allergens, and inflammatory mediators to access the subepithelium [16]. Clinically, barrier dysfunction is a hallmark of asthma, CF, COPD, and acute infections, where junctional breakdown precedes inflammation, neutrophil infiltration, and loss of epithelial homeostasis [17]. It is also implicated in epithelial–mesenchymal transition and immune evasion in lung cancer [17]. Functional assays such as TEER or FITC–dextran flux provide practical correlates of this integrity, while immunostaining for junctional proteins directly reflects structural changes observed in patient biopsies [18]. Measuring barrier function *in vitro,* therefore, captures a disease-relevant phenotype that bridges molecular perturbation with clinically observable pathology, making it a powerful and translatable readout. Here, the baseline system, exposed to the PEG and DEX phases in the absence of bacteria, showed low transepithelial permeability and preserved E-cadherin/ZO-1 architecture, indicating that the ATPS formulation itself does not degrade epithelial integrity and that the model retains dynamic range to resolve injury without saturating the assay. The permeability assay employed standardized macromolecular flux across multiple time points with appropriate diffusion controls, and junctional integrity was corroborated by immunostaining and quantitative network metrics, establishing construct validity for barrier measurements in this platform.

Across biological replicates, PA01 alone increased dextran permeability, indicating barrier compromise; co-culture with *R. mucilaginosa* or *L. casei* significantly reduced PA01-normalized damage to ∼50-60% of PA01 alone, whereas PA01 co-cultured with *S. pneumoniae* produced the highest permeability, consistent with supra-additivity (Figure 4A-D). These phenotypes are consistent with PA01 virulence programs that acutely depress TEER and remodel junctions, LasB and other exoproteins degrade/redistribute tight junction components (ZO-1, occluding, claudins) [19][20], rhamnolipids impair tight junction assembly and promote paracellular transit [21], and T3SS effectors (ExoS/ExoT) reorganize the actin junctional scaffold to enable transmigration [22], which would lead to the permeability surge we observed [23]. Consistent loss of E-cadherin/ZO-1 architecture supports junctional remodelling rather than complete cytolysis, and Live/Dead results exclude group-specific cytotoxic collapse as the likely cause.

The commensal mitigation observed in our model aligns with reports that lactobacilli blunt PA01 driven airway injury *in vivo* [24][25] and that *Rothia* derived metabolites can reprogram *P. aeruginosa* metabolism and attenuate virulence outputs [26], these mechanisms offer plausible routes for the ∼50% rescue in barrier integrity observed here. Conversely, the trend towards synergy with *S. pneumoniae* mirrors clinical and experimental observations that mixed pathogenic communities exacerbate respiratory injury and inflammation [27][28]. Across mono- and polymicrobial conditions, PA01 growth expanded making commensal killing an unlikely explanation for barrier rescue. Cell viability was indistinguishable between conditions, further excluding cytotoxicity as potential effect confounders. These data establish that the airway model we developed detects pathogen mediated barrier failure, resolves commensal protection, and reveals mixed-pathogen potentiation within a single assayable platform.

Cytokine profiling demonstrated feasibility for molecular readouts although results were differential among conditions, modest trends were discernable. The IL-8 pattern of *S. pneumoniae* having the highest expression, is consistent with *S. pneumonia’*s capacity to elicit neutrophil-recruiting chemokines from airway epithelium [31][32]. Co-culture of PA01 with *S. pneumoniae* tending higher than PA01 alone directionally aligns with the supra-additive barrier injury observed functionally. By contrast, PA01’s muted or downward shifts in IL-6/8 relative to the control and to commensals is plausible given known epithelial immunomodulatory by PA01 (e.g., virulence factor mediated interference with junctional and NF-kB signalling, or protease effects on cytokine pathways) [33][34], and may also reflect kinetics, a 24h endpoint can miss early cytokine peaks. Commensals alone elevating Il-6 (and not reducing IL-8) is compatible with transient epithelial sensing that preserves barrier protective signalling without excess inflammation as seen with other commensal taxa [35]. Importantly, CFU data indicate robust PA01 expansion across mono and polymicrobial conditions, arguing against commensal killing as an explanation for lower cytokine output in PA01 groups, viability was comparable across conditions, again, reducing the likelihood that differential cytotoxicity confounded cytokine readouts.

Although underpowered for significance testing, these trends add another layer of fidelity to the platform, complimenting the structural (junctional IF) and functional (permeability) endpoints, cytokine measurements capture condition-specific immunologic tone over the same cultures. Practically, increased early time point (≤6h), basolateral sampling, and multiplex panels coupled with mixed effect modelling could improve inference.

At systems level, the model recapitulates three clinically relevant behaviors: a pathogenic driver (*P. aeruginosa*) that compromises barrier integrity, commensal attenuation (*R. mucilaginosa*, *L. casei*), consistent with protective roles, and supraadditive injury in mixed-pathogen challenge (PAO1 + *S. pneumoniae*). This ‘validation by recapitulation’ confers phenotype and construct validity, the platform reproduces the expected directionality of polymicrobial interactions under controlled conditions. Because it does so with day-scale stability and quantifiable effect sizes, it also provides predictive power that the assay can be used to probe unknown or poorly characterized interactions. In practice, the features that stabilize and facilitate the co-culture (spatial confinement, serial readouts) overcome the resolution and collapse constraints that have previously limited discovery in traditional culture systems, enabling systematic hypothesis generation about emergent polymicrobial effects.

Because these phenotypes are revealed in a format that accommodates serial permeability assessment, junctional imaging, and soluble mediator sampling, the platform resolves interaction-dependent effects that often remain obscured in traditional culture setups. The approach is therefore ideal for mechanistic studies of diseases where dysbiosis and barrier failure co-evolve, such as cystic fibrosis, COPD, and bronchiectasis, and for emerging questions at the interface of tumor biology, EMT, and immune evasion in lung cancer, where epithelial permeability and junctional remodeling are central axes of pathophysiology.

## Limitations

Our model is reductionistic by design, reflecting an intended scope of use (i.e., controlled, day scale investigation of polymicrobial interaction dependent epithelial phenotypes rather than a full recapitulation of the airway niche). However, limitations of our model include; first, the cellular composition is limited to an epithelial-endothelial set up without professional immune cells; consequently, neutrophil or macrophage mediated junctional damage, immune mediated cytotoxicity, and cytokine amplification loops are not captured. Similarly, the microbial communities are simplified relative to the *in vivo* ecosystem, higher-order interactions (e.g., keystone metabolites, cross-feeding, viral or fungal interactions) are likely under-represented. This could be addressed by introducing patent derived multitaxon (microbiome) samples to assess model performance and validation across microbial variability. Methodologically, the permeability assay captures net apical-basal macromolecular transport but does not resolve the specific route (paracellular vs transcytotic) or mode (short lived pore states or sustained junctional remodelling) and spatially averages heterogeneity across the whole insert. Pairing FITC-dextran with TEER and additional junctional imaging could help distinguish ionic from macromolecular pathways and localize leaks. Finally, between run dispersion in cytokines reduced power to detect subtle effects, a limitation mitigatable by increasing biological replicates, internal controls, mixed effects modelling that accounts for run variations, and earlier time point sampling. These constraints limit direct physiological extrapolation but do not detract from the platform’s core strength: controlled, spatially stabilized modelling of interaction-dependent microbe-epithelial phenotypes

## Future directions

The lung airway model is readily extensible and its reductionistic design reflects its intended scope of use, this however is intended to be adaptable for a variety of investigations. Adding an immune component (e.g., neutrophils, macrophages, or NK cells introduced in a time-staggered manner to preserve epithelial viability) would enable investigation of host amplification and response phases, including effector impact on junctional dynamics. Expanding to more complex, patient-derived microbial samples will permit mapping of higher-order interactions and identification of community features that predict barrier outcomes. Methodologically, pairing TEER with FITC-dextran will help partition ionic versus macromolecular pathways, automated live imaging could quantify barrier failure kinetics. Finally, factorial assessment integrated with multi-omics isolating pathogen virulence (T3SS mutants, elastase inhibition, quorum sensing blockers) from commensal signals (purified post biotics) could help dissect mechanism of action for the emergent effects.

## Conclusion

By enforcing spatial confinement of microbial growth, ATPS converts a fragile, short-lived co-culture into a robust, assayable platform that preserves epithelial function long enough to expose interaction-dependent phenotypes. The model both reproduces established behaviors (pathogen injury, commensal mitigation, mixed-pathogen synergy) and enables multi-modal, longitudinal measurements that are generally inaccessible in conventional *in vitro* systems. Together, these findings highlight the need of interaction-aware models and establish ATPS-confined airway cultures as an assayable, physiologically relevant platform for dissecting host–microbiome dynamics and for accelerating mechanism guided therapeutic discovery.

## Author contributions

BL and SS conceptualized the project. SS designed and acquired the experiments with assistance from BL and KV. SS and KV wrote the manuscript. All authors reviewed the manuscript, gave feedback and have approved the final version of the manuscript.

## Acknowledgements

The authors gratefully acknowledge Dr. Zenyu Cheng and Dr. Jason LeBlanc for their contributions of bacterial strains. This work was supported through funding provided by the Natural Sciences and Engineering Research Council of Canada (NSERC) Discovery Grant program (RGPIN-2018-05742) and the Canada Foundation for Innovation − John R. Evan Leaders Fund (project# 36032). SS was a recipient of the Canada Graduate Research Scholarship (Doctoral).

